# Wild rice *Oryza rufipogon* outperforms cultivated rice in stimulating beneficial bacterial endophytes

**DOI:** 10.64898/2026.05.22.727159

**Authors:** Francesca Vaccaro, Maria Laura Amenta, Iacopo Passeri, Camilla Fagorzi, Stefano Varriale, Danuše Tarkowská, Aleš Pěnčík, Ivan Petřík, Federica Brunoni, Vittoria Brambilla, Andrea Rossoni, Erica Mica, Giampiero Valè, Elena Perrin, Alessio Mengoni, Roberto Defez, Carmelina Bianco

## Abstract

Beneficial interactions between plants and microorganisms strongly influence plant health and productivity, and root exudates play a central role in shaping these associations. This study analyzed the transcriptional responses of the bacterial endophytes *Enterobacter asburiae* RCA24 and *Kosakonia sacchari* RCA25 to root exudates from two commercial Italian rice accessions (*Oryza sativa* Baldo and Vialone Nano) and from an accession of the wild progenitor of tropical rice, *Oryza rufipogon*. Bacterial transcriptome analyses revealed that RCA24 responds differently to *O. sativa* varieties and that RCA25 was more stimulated by *O. rufipogon*. Changes in bacterial gene expression were mainly related to central metabolism, stress response, and signal transduction, highlighting a precise pattern of interaction. On the other hand, transcriptome analysis of inoculated rice revealed that RCA24 triggered broader transcriptional changes in plants than RCA25. Differentially expressed genes were related, especially in shoots, to defense responses, hormone-mediated signaling, and ribosome biogenesis, revealing that plants discriminate bacterial strains in a genotype-specific manner at the transcriptional level. Our findings suggest that traits beneficial to plant-soil microbiota interactions present in *O. rufipogon* and lost during domestication and diversification could be identified and reintroduced into modern rice varieties to improve sustainable field performance through beneficial microbial associations.

## Introduction

Plant-microbe interactions are fundamental for plant health and productivity (Hassani *et al*., 2018). Crop yield and resilience are influenced by multiple factors, including plant genotype, agronomic practices, the environment, and the crop’s associated microbiome (Berg, 2009; Vandenkoornhuyse *et al*., 2015). Identifying plant genotypes more tolerant to abiotic and biotic stresses, highly productive, and capable of benefiting further from positive interactions with microorganisms is central to what is known as microbiome-driven or microbiome-assisted plant breeding (Nerva *et al*., 2022). Microorganisms can provide plants with the molecules necessary for growth, which are usually supplemented externally through fertilization, and can protect against pathogens’ attack. Efficient plant-microbial symbiont associations rely on complex interactions between the genotypes of both partners, which are still poorly understood. Rice is the first crop consumed by humans worldwide, and it is harvested on 160-170 million ha (FAOSTAT, 2024), second only to maize. Italy is the leading European rice producer, with 226.000 ha cultivated in 2024 (Ente Nazionale Risi), which represents 50-55% of the total rice cultivated area in Europe (FAOSTAT, 2024). Cultivated rice belongs to the O*ryza sativa* species, which was domesticated from its wild progenitor, *O. rufipogon*, an endemic species to the wetlands of tropical and subtropical Asia (Gutaker *et al*., 2020). Italian rice cultivars, including the commercial ones used in this study, “Baldo” and “Vialone nano”, generally belong to the temperate *japonica* group, which separated from *indica* after domestication and diversified on independent trajectories. *O. rufipogon* possesses greater allelic variation than domesticated *O. sativa*, and it is a donor for most of the genetic variability that has been lost during the domestication and breeding of rice (Izawa, 2022). Domestication has influenced the composition of the root-associated microbiome (Escudero-Martinez and Bulgarelli, 2019; Pérez-Jaramillo *et al*., 2018; Raaijmakers and Kiers, 2022). Although genotypic differences in rice have been observed to impact root-associated microbial communities significantly (Edwards *et al*., 2015; Xiong *et al*., 2021; Xu *et al*., 2020), rice breeding programs have so far not paid sufficient attention to analyzing the root-associated microbiome. Root exudates mediate the first contact between roots and the soil microbiota (Korenblum *et al*., 2020). Root exudates are a diverse group of molecules secreted by plants into the rhizosphere, the soil area immediately surrounding the roots. These exudates can serve as a source of nutrients for microbes and as signaling molecules that can influence microbial behavior and gene expression (Korenblum *et al*., 2020; Zhalnina *et al*., 2018; Poole *et al*., 2018). Different host plant genotypes produce root exudates with distinct chemical compositions, which give rise to distinct symbiotic interactions (Kiers *et al*., 2007; Pacheco-Moreno *et al*., 2024; Weese *et al*., 2015).

By studying the transcriptomic response of the rhizobium *Sinorhizobium meliloti* to root exudates of different *Medicago sativa* (alfalfa) cultivars, it was demonstrated that nearly 30% of the total differentially expressed genes could be statistically correlated with the interaction between specific plant and bacterial genotypes (Fagorzi *et al*., 2021). Similar evidence has been recently demonstrated for the transcriptomic response between Bacteria and Fungi, belonging to the plant microbiota (Vaccaro *et al*., 2024), suggesting that specificity of genotype-by-genotype interactions is a key determinant of the successful assembly of a plant microbiota. We speculate that a parallel mechanism might occur in endophyte-rice communications. Whole transcriptome analysis used to uncover the rice molecular networks during symbiotic relationships highlighted several actors, including genes related to secondary metabolism, immunity, and phytohormones (King *et al*., 2023), as well as ammonium and sugar transport (Thomas *et al*., 2019).

To investigate the differential endophytic colonization ability of rice genotypes, we analyzed the differential transcriptomic response in the early phase of plant root colonization, corresponding to the perception of root exudates. To test this hypothesis, we used an experimental model consisting of two endophytic plant growth promoting strains (*E. asburiae* RCA24 and *K. sacchari* RCA25), which had a different ability to colonize the wild specie *O. rufipogon* accession 602-131-2 compared to the *O. sativa* varieties ‘Vialone Nano’ and ‘Baldo’ (Andreozzi *et al*., 2019), and three plant genotypes corresponding to two commercial varieties of temperate Italian rice (*O. sativa*) and one accession of the tropical wild ancestor *O. rufipogon*.

The results indicated that the two bacterial strains could reprogram their transcriptomes in response to the presence of a particular rice cultivar or species, both in terms of the number of genes affected (ranging from 0.1 to 20% of total expressed genes) and the type of gene functions. The wild *O. rufipogon* accession affected a greater number of signal transduction-related functions than the cultivated ones. We also verified that the three rice genotypes released distinct phytohormone profiles in the rhizosphere, with greater genotypic differentiation for gibberellins than for auxins. Our results highlight a reciprocal dialogue between plants and bacteria: the wild ancestor *O. rufipogon* not only “communicates” differently with RCA25 *via* exudates, but also “listens” and responds by activating specific gene pathways in its roots.

## Materials and methods

### Bacterial strains and culture conditions

*E. asburiae* RCA24 and *K. sacchari* RCA25 were originally isolated from *O. sativa* temperate *japonica* cv. Volano rice plants grown in Italy (Defez *et al*., 2017). *E. asburiae* RCA24 was shown to produce indole-3-acetic acid, while *K. sacchari* RCA25 harbors nitrogen-fixation genes and fixes atmospheric nitrogen inside rice plants (Bianco *et al*., 2021). Strains were grown at 30 °C in LB medium containing 20 µg ml^-1^ vancomycin. Thereafter, the two strains are indicated as RCA24 and RCA25 only.

Genome sequences and annotations of RCA24 and RCA25 strains can be accessed from Bioproject PRJNA1120238 (assemblies ASM4012233v1 and ASM4012231v1, respectively).

### Plant materials

Three rice accessions were examined in this study: *O. sativa* ‘Vialone Nano’ and *O. sativa* ‘Baldo’, which are commercial varieties of Italian rice within the temperate *Japonica* group, and *O. rufipogon* accession 602-131-2, (kindly provided by IRRI, International Rice Research Institute, Metro Manila, Philippines).

### Rice colonization efficiency

Rice seeds were surface sterilized by incubation in 70% ethanol for 8 min, followed by 5% sodium hypochlorite solution containing 0.1% Tween 20 for 25 min. Seeds were then washed several times with sterilized distilled water and incubated for five days at 21 °C in the dark for germination. Germinated seeds were transferred into plastic pots containing inorganic substrates (sand with a 1.2–0.8 mm granule size and perlite with a 3–4 mm granule size) in a 1:1 ratio. Each planting unit was kept in the growth chamber under long-day conditions (16 h), at 19–23 °C, and 75% relative humidity. 21 days after inoculation (DAI), plants were carefully uprooted from pots, washed by dipping them in tap water, and weighted. Surface sterilization of whole plants and tissue homogenization were carried out as described in Bianco *et al*. (2021). Dilutions of the homogenates (1:10,000) were spread onto 1.5% LB agar plates containing the specific antibiotic (20 μg mL^-1^ vancomycin) and incubated at 30 °C. After 24 h, CFU were counted.

### Seeds inoculation and plant growth

Dehulled seeds of rice plants (‘Baldo’, ‘V. Nano’, and ‘Rufipogon’) were surface-sterilized and germinated as described by Amenta *et al*. (2024). After 5 days, germinated seeds were incubated in Petri dishes with 50 mL of 1x PBS solution containing each strain to a final concentration of 10^7^ cells mL^−1^ for 4 h at room temperature. Seeds incubated in 1x PBS were used as a control. Inoculated seeds were then transferred into plastic pots containing a 1:1 mixture of sand (0.8–1.2 mm granule size) and perlite (3–4 mm granule size) as a substrate. Each planting unit was kept in the growth chamber under long-day (16 h) conditions, at 19°C and 75% relative humidity, and watered daily. Once a week, a nitrogen-free nutrient medium (Bianco *et al.,* 2021) was added to the plants. Six replicates were set up for each rice ecotype.

### Root exudates preparation and bacterial treatment

For exudate collection, the roots of non-inoculated, intact (three-week-old) plants were carefully removed from pots, washed to remove any remaining soil, transferred to 30 mL Milli-Q water, and incubated for 48 h in greenhouse conditions. After 48 h of incubation, each root exudate solution was filtered through a 0.22 μm Millipore filter membrane to remove any root debris, cold freeze-dried, and then stored at −80 °C until further hormonal analysis. Aliquots of non-dehydrated samples were used to treat RCA24 and RCA25 bacterial cultures.

Overnight cultures of both strains were diluted with fresh LB medium. When the OD_600_ reached 0.6, they were supplemented with 0.5 ml of root exudates collected from rice plants. The same amount of Mili-Q water served as a control. The volume of root exudates for bacterial treatment was selected, since no discernible effect on bacterial growth was observed at this level. The cultures were grown at 30°C and 200 rpm for 2.5 h, and cells were harvested for RNA isolation. For each strain and treatment, three biological replicate cultures were prepared, yielding a total of 12 samples for strain.

### Detection and quantification of phytohormones in root exudates

The sample preparation and analysis of gibberellins (GAs) were performed according to the method described in Urbanová *et al*. (2013) with some modifications. Briefly, rice root exudate samples evaporated to dryness were dissolved in 1 mL of ice-cold 80 % acetonitrile containing 5 % formic acid as the extraction solution, and then sonicated for 5 min in a laboratory ultrasonic bath. The samples were then extracted overnight at 4 °C using a benchtop laboratory rotator Stuart SB3 (Bibby Scientific Ltd., UK) after adding 2 pmol of [^2^H_2_]GA_1_, [^2^H_2_]GA_4_, [^2^H_2_]GA_9_, [^2^H_2_]GA_19_, [^2^H_2_]GA_20_, [^2^H_2_]GA_24_, [^2^H_2_]GA_29_, [^2^H_2_]GA_34_ and [^2^H_2_]GA_44_ (OlChemIm, Czech Republic) as internal standards. The homogenates were centrifuged at 36,670 *g* and 4 °C for 10 min (Hermle Z 35 HK, Hermle Labortechnik GmbH, Germany), and corresponding supernatants were further purified using mixed-mode SPE cartridges (Waters, Ireland). After SPE purification, the samples were evaporated to dryness in a stream of nitrogen (Turbovap® LV, Biotage, Sweden), redissolved in the starting mobile phase, and analyzed by ultra-high performance liquid chromatography-tandem mass spectrometry (UHPLC-MS/MS; Waters, USA) using an Acquity UPLC I-Class Plus system (Waters, USA) coupled to a triple quadrupole mass spectrometer Xevo TQ-XS (Waters, USA). GAs were detected using the multiple-reaction monitoring mode, monitoring the transition of the [M–H]-ion to the appropriate product ion, as described in Urbanová *et al*. (2013). Masslynx 4.2 software (Waters, USA) was used to analyze the data. GA levels were quantified using the standard isotope dilution method (Rittenberg and Foster, 1940).

Auxin metabolites were quantified as described by Pěnčík *et al*. (2018). Dried root exudate samples were dissolved in 1 mL 50 mmol/L phosphate buffer (pH 7.0) containing 0.1% sodium diethyldithiocarbamate and stable isotope-labeled internal standards. Aliquots of 200 µL were either acidified to pH 2.7 and purified by in-tip µSPE, or derivatized with cysteamine prior to acidification and µSPE for IPyA determination. Eluates were evaporated, reconstituted in 10% aqueous methanol, and analyzed using an HPLC system 1290 Infinity III (Agilent Technologies, USA) equipped with Kinetex C18 column (50 mmx2.1 mm, 1.7 µm; Phenomenex) and linked to 6495D Triple Quad detector (Agilent Technologies, USA).

Phytohormone concentrations were log□□(x+1)-transformed prior to multivariate analysis to reduce the influence of extreme values and homogenize variance. Variables with zero variance across samples were excluded. Principal Component Analysis (PCA) was performed on centered, non-scaled data using the *prcomp* function in R, and results were visualized as biplots using the *ggplot2* and *ggfortify* packages. Differences in phytohormone profiles among genotypes were assessed by permutational multivariate analysis of variance (PERMANOVA) using the *adonis2* function from the *vegan* package, with genotype as the explanatory variable and 999 permutations. Mean concentrations ± standard deviations were calculated for each compound and genotype from three biological replicates. All analyses were performed in R (v.4.4.2).

### RNA preparation for RNA-seq analysis

Bacterial cultures of RCA24 and RCA25 were grown in LB medium to the exponential phase (OD_600_ = 0.7) and then treated with root exudates of *O. sativa* ‘Vialone Nano’, *O. sativa* ‘Baldo’ and *O. rufipogon.* Untreated cultures were used as controls. RNA was extracted from RCA24 and RCA25 using the PureLink RNA Mini Kit (Invitrogen) following the manufacturer’s instructions, with the following modification: proteinase K at 75 µg ml^-1^ was added to the lysis buffer. Residual DNA present in the RNA preparations was removed using the RNase-free TURBO DNase kit (Ambion) according to the manufacturer’s instructions. Quality of RNA was checked by measuring the RNA Integrity Number (R.I.N.) on TapeStation automated electrophoresis apparatus (Agilent, CA, USA).

For plant RNA preparation, roots and leaves from two-week-old non-inoculated and RCA24- and RCA25-inoculated plants were collected, frozen in liquid nitrogen, and stored at −80 °C until use. Total RNA was extracted from frozen material (100 mg) using the Trizol Reagent (Invitrogen) according to the manufacturer’s instructions. Residual DNA present in the RNA preparations was removed using the RNase-free TURBO DNase kit (Ambion) according to the manufacturer’s instructions. The quality of the RNA was assessed by measuring the R.I.N. using the Agilent 2100 Bioanalyzer and the Caliper GX.

### RNA sequencing and analysis

A bacterial RNA sequencing library was constructed after rRNA depletion using the RiboZero kit (Illumina) with the TruSeq Stranded Total RNA kit (Illumina). Sequencing was performed by IGA Technology Service (Udine, Italy). Plant RNA sequencing library preparation and sequencing was performed by BMKGENE technology service (Münster, Germany).

Raw bacterial RNA-seq reads were subjected to quality control and preprocessed using fastp (v0.23.4) to ensure high-quality data for downstream analysis. fastp was used with default settings. The quality of reads before and after preprocessing was assessed using FastQC (v0.12.1). A reference-guided transcripts assembly was then performed. All reads datasets were aligned using HISAT2 (v2.2.1) with default parameters over the indexed genomes (Kim *et al*., 2019). The sequence alignment mapping results collected have been compressed to BAM format, sorted, indexed, and merged using samtools (Li *et al*., 2009). The assemblies were run using StringTie2 (v2.2.3) (Kovaka *et al*., 2019) with the “-G” option to include the annotation in the merging (refer to “Annotation” paragraph for more details) and to obtain the GTF of covered transcripts. The transcriptome in FASTA format was then extracted by cross-referencing the GTF and the reference genome using gffread (v0.12.8) (Pertea and Pertea, 2020). The genome annotations were performed using Prokka (1.14.6) with default options (Seemann, 2014). Functional enrichment was obtained using eggNOG mapper across the entire gene set, annotating for GOs covering molecular functions, KEGG pathways, biological processes, and cellular components. To retrieve the aminoacidic sequences required by eggNOG mapper, the Seqtk (v1.4) tool has been used with a list of the collected coding sequences (CDS). Salmon (v1.10.3) (Patro *et al*., 2017) was used to estimate the transcripts abundances. After the transcriptome indexing, quantification was obtained by running Salmon with the “--validateMappings” option for selective alignment. Differential abundance analysis was performed with the DESeq2 version 1.36.0 package (Love *et al*., 2014). Differentially expressed genes (DEGs, log_2_FC threshold > 1, Padj *<*0.05) of RCA24 and RCA25 were further functionally annotated using the online version of eggNOG-mapper version 2 with default settings (Cantalapiedra *et al*., 2021), which assigns KEGG Orthology (KO) identifiers based on sequence homology against the eggNOG database. The resulting KO assignments were used to map DEGs to metabolic pathways in the KEGG database. For each pairwise contrast, the number of DEGs mapping to each KEGG pathway was counted using the KEGG_Pathway field from the eggNOG-mapper output, retaining only entries with a valid ‘map’-prefixed pathway identifier. Pathway names were retrieved programmatically via the KEGGREST R package (Tenenbaum, 2025). All statistical analyses were performed in R Studio (ver. 4.2.0) (RStudio Team, 2020). ANOVA and Tukey’s tests were performed using *aov* and *TukeyHSD* functions, respectively. Heatmaps were generated using the *pheatmap* package. PCA was performed using the *prcomp* function of R, and the results were visualized with the *ggplot2* package (Wickham, 2009). Venn diagrams were obtained with *ggvenn* package.

Raw plant RNA-seq reads were preprocessed and cleaned by removal of adaptor sequences, low-quality-end trimming, and low-quality reads using fastp v1.0.1 with default settings. Transcript-level quantification was performed with Salmon v1.10.1 (Patro *et al*., 2017) against the CDS reference transcriptome deposited on Ensembl Plants to restrict quantification to protein-coding regions. The quantification results were collected with custom scripts. The resulting gene-counts matrix was imported into DESeq2 (Love *et al*., 2014) for downstream analysis. Genes were filtered for padj < 0.05 and log2FC > 1. Data analysis and visualization were performed in RStudio v5.4.1. Gene annotations were retrieved from Ensembl Plants and intersected with the set of DEGs for functional characterization. To identify significantly overrepresented functional categories within DEGs, enrichment analysis was performed using COG categories via the eggNOG-mapper web server (version 2) with default parameters. COG categories provided a compact, non-redundant categorization scheme that facilitated visualization of general enriched functions within DEGs. Nested likelihood ratio tests (LRTs) were performed as indicated previously (Fagorzi *et al*., 2021). The DEGs identified as significant in the interaction model were used to perform Gene Onthology (GO) enrichment analysis with *topGO* (Alexa and Rahnenfuhrer, 2024) to provide a pathway-centred analysis for enrichment relative to those DEGs.

## Results

### E. asburiae RCA24 and K. sacchari RCA25 differently colonized rice plants

Quantitative evaluation of endophytic colonization revealed sharp differences in host preference. RCA25 efficiently colonized *O. rufipogon* and ‘Baldo’ (12.9-fold and 3.2-fold more than RCA24, respectively), while RCA24 showed a 10-fold preference for ‘Vialone Nano’ (Table 1).

**Table 1.** Colonization efficiency of rice plant genotypes by RCA24 and RCA25.

| Rice variety | *E. asburiae* RCA24<br>CFU/g<sup>a</sup> | *K. sacchari* RCA25<br>CFU/g<sup>a</sup> | Ratio<br>RCA25 vs. RCA24 |
| --- | --- | --- | --- |
| *O. sativa* Baldo | 6.8 ± 1.3 x10<sup>8</sup> | 22.0 ± 3.7 x10<sup>8</sup> | 3.2 ±0.8 |
| *O. sativa* Vialone Nano | 24.5 ± 4.1 x10<sup>8</sup> | 2.4 ± 0.6 x10<sup>8</sup> | 0.10 ±0.03 |
| *O. rufipogon* | 9.5 ± 1.0 x10<sup>8</sup> | 123 ± 15 x10<sup>8</sup> | 12.9 ±2.1 |

<sup>a</sup>Mean and standard deviation of colonization efficiency (CFU/g fresh weight) for six independent biological replicates.

### Differentiated transcriptional response of E. asburiae RCA24 and K. sacchari RCA25 to rice root exudates

Transcriptome analysis of RCA24 and RCA25 exposed to root exudates demonstrated precise reprogramming. A relevant result from the analysis of the differentially expressed genes (DEGs) (Dataset S1) indicated that *O. rufipogon* exudates elicited nearly 6 times more DEGs in RCA25 than in RCA24 (Table 2). Principal Component Analysis (PCA) showed that bacterial responses were highly genotype-specific (Fig. 1). When the transcriptomes of RCA24 and RCA25 were compared (Fig. 2), we found that the treatment with ‘Vialone Nano’ root exudates led to the 66.7 % of the identified DEGs in RCA24, while the treatment with *O. rufipogon* root exudates produced the highest number of DEGs (93.7% of the total) in RCA25. By contrast, the treatment with ‘Vialone Nano’ exudates resulted in the fewest unique DEGs in RCA25, accounting for only 1.3% of the total DEGs. Concerning the specific functional features of the affected transcriptomes, the overall DEGs were represented by different functions depending on strain (RCA24 or RCA25) and contrast (*O. sativa* and *O. rufipogon* vs control or *O. sativa* vs *O. rufipogon* root exudates) (Fig. 3, Table S1). The Clusters of Orthologous Groups (COG) S category (Function unknown) was abundantly present in all datasets, indicating that gene functions involved in bacteria-plant recognition are still poorly characterized. In general, categories related to central metabolism ([E] amino acid transport and metabolism; [C] energy production and conversion; [G] carbohydrate transport and metabolism; [H] coenzyme transport and metabolism) were present in almost all contrasts, either among up- and down-regulated DEGs (Dataset S1). Nearly all the contrasts with *O. rufipogon* produced DEGs mapping on the two-component system of signal transduction, while the same pathway was identified only for RCA24 treated with ‘Vialone Nano’ vs control. We may speculate that *O. rufipogon* induced greater remodeling of cell behavior compared to *O. sativa* varieties’ root exudates. Nitrogen metabolism was downregulated in RCA25 in both ‘Baldo’ and ‘Vialone Nano’ vs *O. rufipogon* comparisons and upregulated in RCA24 exposed to ‘Vialone Nano’ and *O. rufipogon* exudates vs control. Cysteine and methionine metabolism was found exclusively in RCA24, appearing in multiple contrasts, while it was completely absent in RCA25. Regarding specific genes, iron transport (*feoA*) was upregulated in both RCA24 and RCA25 in response to *O. rufipogon* exudates vs control, representing one of the few convergent responses between the two strains. Hydrogenase assembly (*hypA*) showed a more complex pattern, being downregulated in RCA24 with Baldo exudates and upregulated in RCA25 with *O. rufipogon* exudates, suggesting strain-specific metabolic adjustments. Purine biosynthesis (*purK*) was regulated in multiple contrasts in both strains. It was downregulated in RCA24 exposed to ‘Vialone Nano’ exudates and in RCA25 exposed to *O. rufipogon* exudates and upregulated in RCA25 with ‘Baldo’ exudates. An opposite amino acid metabolism was observed between the two strains in the presence of wild rice exudates: aromatic amino acid biosynthesis (*aroG*) was reduced in strain RCA25 exposed to *O. rufipogon* exudates compared to the control, while methionine biosynthesis (*metB*) was increased in strain RCA24 under the same conditions. Among genes with strain-specific regulation, the putative selenoprotein *ydfZ* and the peptidase *yhbU* were upregulated in RCA24 exposed to ‘Vialone Nano’ exudates vs control but downregulated in RCA25 in contrasts involving *O. rufipogon*. The nickel/cobalt stress response gene *yfgG* was consistently upregulated in RCA24 in contrasts involving Vialone Nano exudates.

**Fig. 1.**
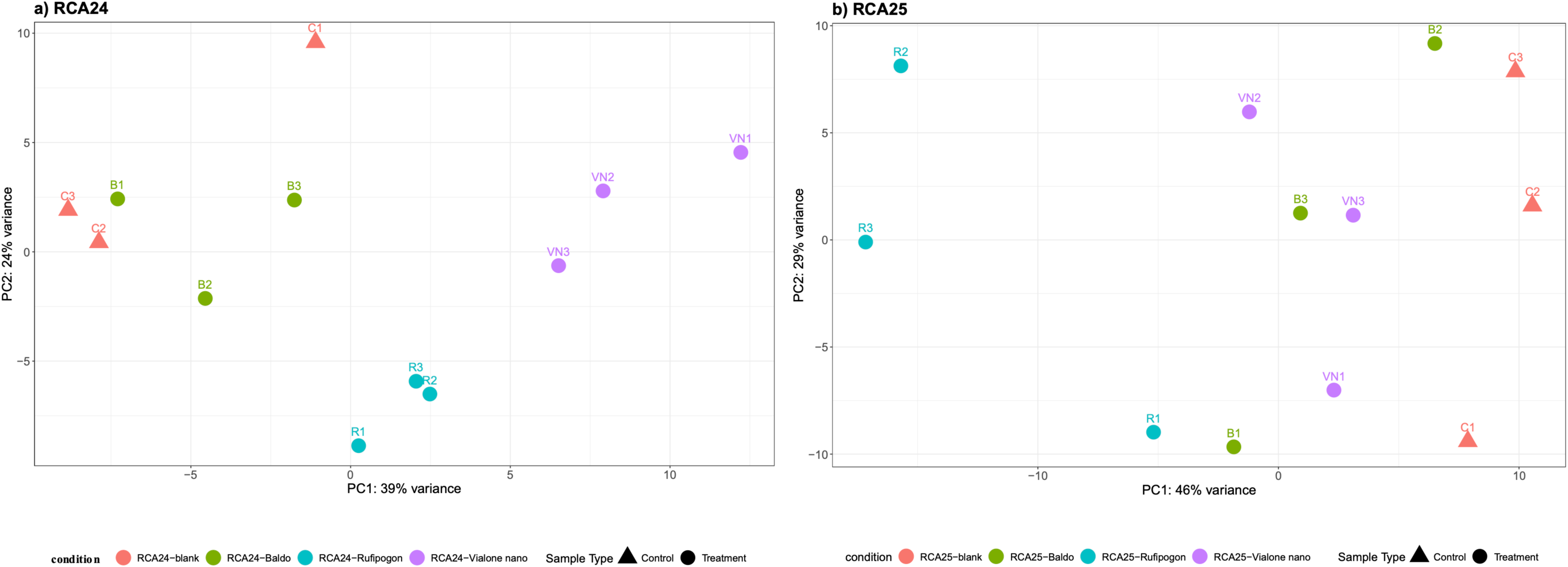
Principal Component Analysis of expression values before filtering by p-value across replicates. Overall differences in DEGs elicited by root exudates of *O. sativa* and *O. rufipogon* in A) RCA24 and B) RCA25. The centroids refer to single replicate. B, Baldo; R, Rufipogon; VN, Vialone Nano; C, control.

**Fig. 2.**
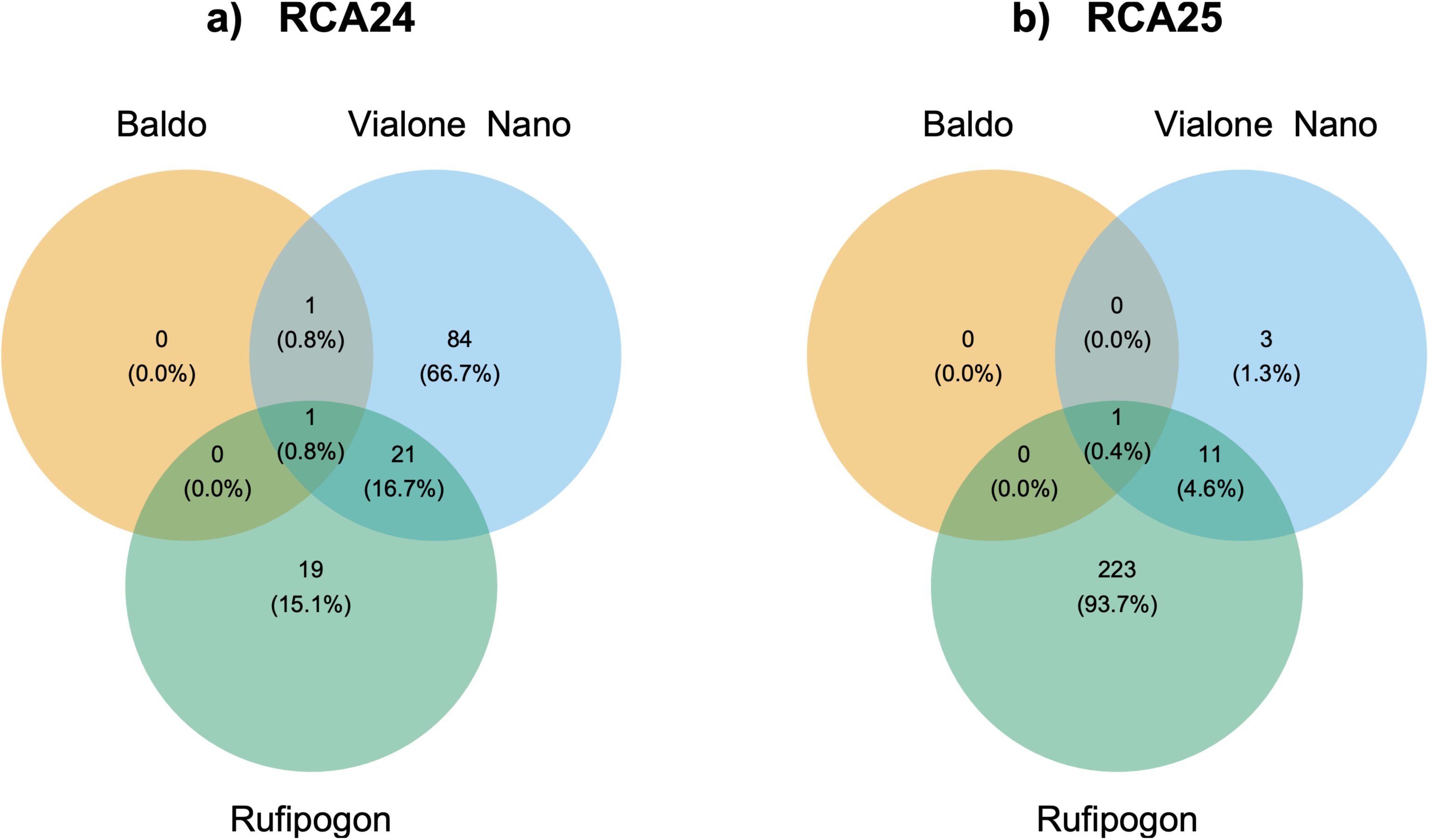
Venn diagram of unique and shared DEGs for RCA24 and RCA25 transcriptome against control (ctr). Percentages are relative to the overall DEGs identified per each strain.

**Fig. 3.**
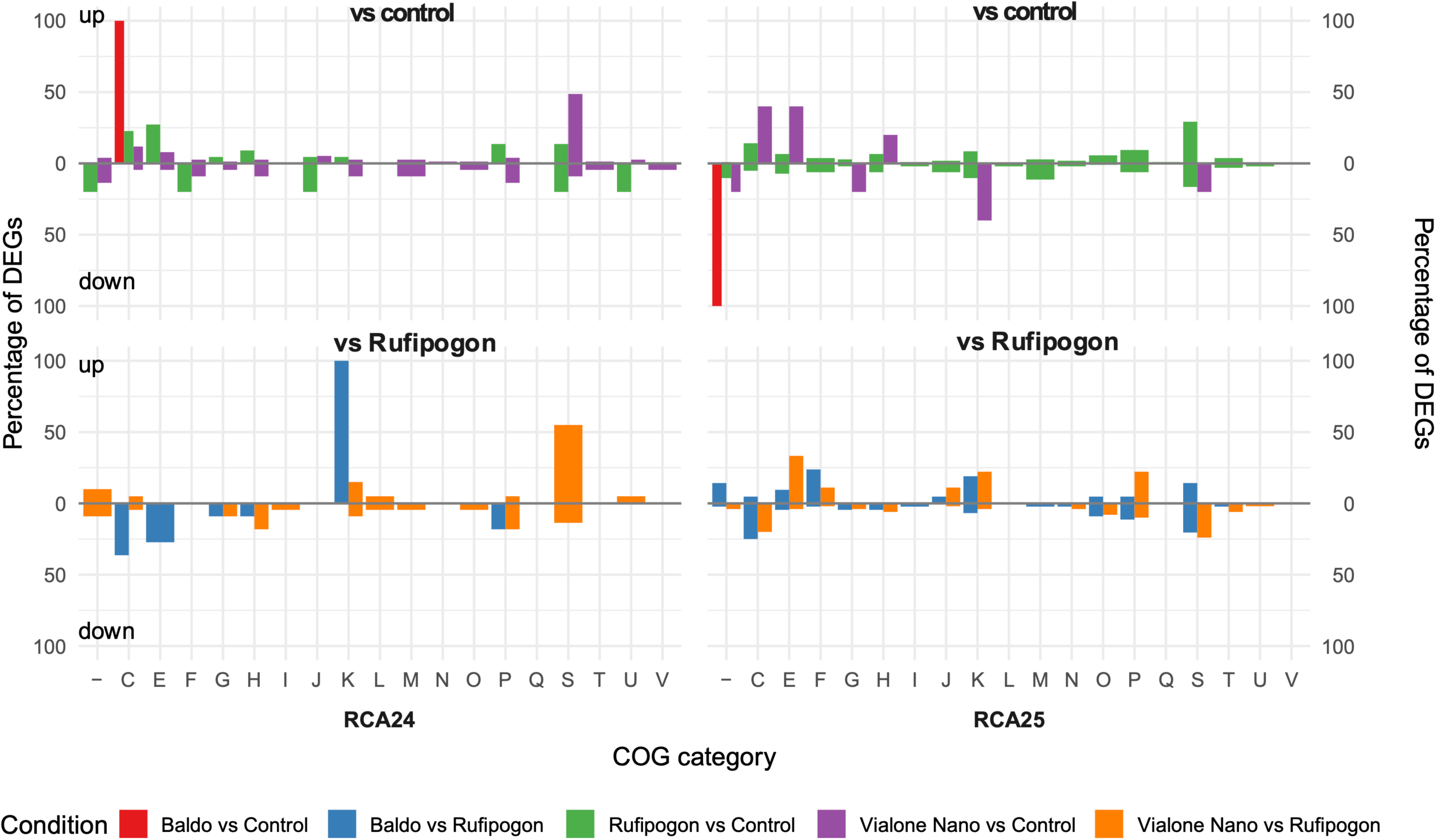
Bar plots showing the percentage of upregulated and downregulated DEGs mapped to COG functions for strains RCA24 (left) and RCA25 (right) compared to control and to *O. rufipogon* (see Table S1 for COG one letter code description).

**Table 2.** Number of DEGs identified in RCA24 and RCA25.

| Contrast | *E. asburiae* RCA24 | *K. sacchari* RCA25 |
| --- | --- | --- |
| Baldo vs. control | 2 (0.05%) | 1 (0.02%) |
| Vialone Nano vs. control | 107 (2.48%) | 15 (0.32%) |
| *O. rufipogon* vs. control | 41 (0.95%) | 235 (5.13%) |
| Baldo vs *O. rufipogon* | 15 (0.35%) | 70 (1.53%) |
| Vialone Nano vs *O. rufipogon* | 57 (1.32%) | 63 (1.37%) |

The percentages in relation to the total number of genes are reported in brackets (genes are 4,309 and 4,575 for RCA24 and RCA25, respectively).

### Phytohormones released into the rhizosphere by three rice genotypes exhibited distinct profiles

Root exudates and root extracts from *O. sativa* (‘Baldo’ and ‘Vialone Nano’) and *O. rufipogon* were evaluated for the presence and abundance (pmol/l) of phytohormones belonging to the classes of auxins and gibberellins, key phytohormones that are widely manipulated by microorganisms to aid colonization and promote growth (Mukherjee *et al*., 2022) (Table S2, Table S3, Table S4, Fig. S1). Gibberellin (GA) and auxin profiles varied significantly across genotypes (Fig. 4A, B). For auxins, exudates containing both active and inactive conjugated forms, PC1 and PC2 explained 50.6% and 22.5% of the variance, respectively (Fig. 4A). PC1 was driven by indole-3-acetic acid (IAA) and tryptamine (TRA), most abundant in Baldo (mean IAA: 33.2 pmol/l; TRA: 2.3 pmol/l), while the inactive form IAA-glutamate (IAA-Glu) was detectable only in ‘Vialone Nano’ (mean: 4.0 pmol/l). PC2 reflected opposing gradients of indole-3-pyruvic acid (IPyA), which were highest in ‘Vialone Nano’ (391.9 pmol/l vs. 179.1 pmol/l in ‘Baldo’ and 158.2 pmol/l in Rufipogon), and 2-oxoindole-3-acetic acid (oxIAA), which increased from ‘Baldo’ (24.4 pmol/l) through Rufipogon (41.8 pmol/l) to ‘Vialone Nano’ (61.4 pmol/l). Genotype explained 39% of the total multivariate variance (PERMANOVA: R² = 0.39, F = 1.96, p = 0.034). For gibberellins, the measured levels ranged by several orders of magnitude, from 0.008 pmol/l (the inactive form GA_34_ in *O. sativa* cv. Baldo) to 956 pmol/l (the inactive form GA_9_ in *O. sativa* cv. Baldo). PC1 explained 77.1% of the variance (Fig. 4B), driven by the precursor GA_9_ (highest in Baldo, mean: 956.4 pmol/l) and the inactive form GA_51_ (highest in Rufipogon, mean: 577.4 pmol/l), with ‘Vialone Nano’ showing intermediate values for both (GA_9_: 219.2; GA_51_: 74.8 pmol/l). Among the active gibberellins (GA_1_, GA_3_, GA_4_, and GA_7_), GA_4_ was the most abundant, with the highest values observed for ‘Vialone Nano’ (mean: 2.82 pmol/l). Remaining gibberellins, including both precursors and inactive forms, were present at < 5 pmol/l. Genotype explained 53% of the multivariate variance (PERMANOVA: R² = 0.53, F = 3.36, p = 0.006). Overall, the three genotypes released distinct phytohormone profiles into the rhizosphere, with stronger genotypic differentiation for gibberellins than auxins.

**Fig. 4.**
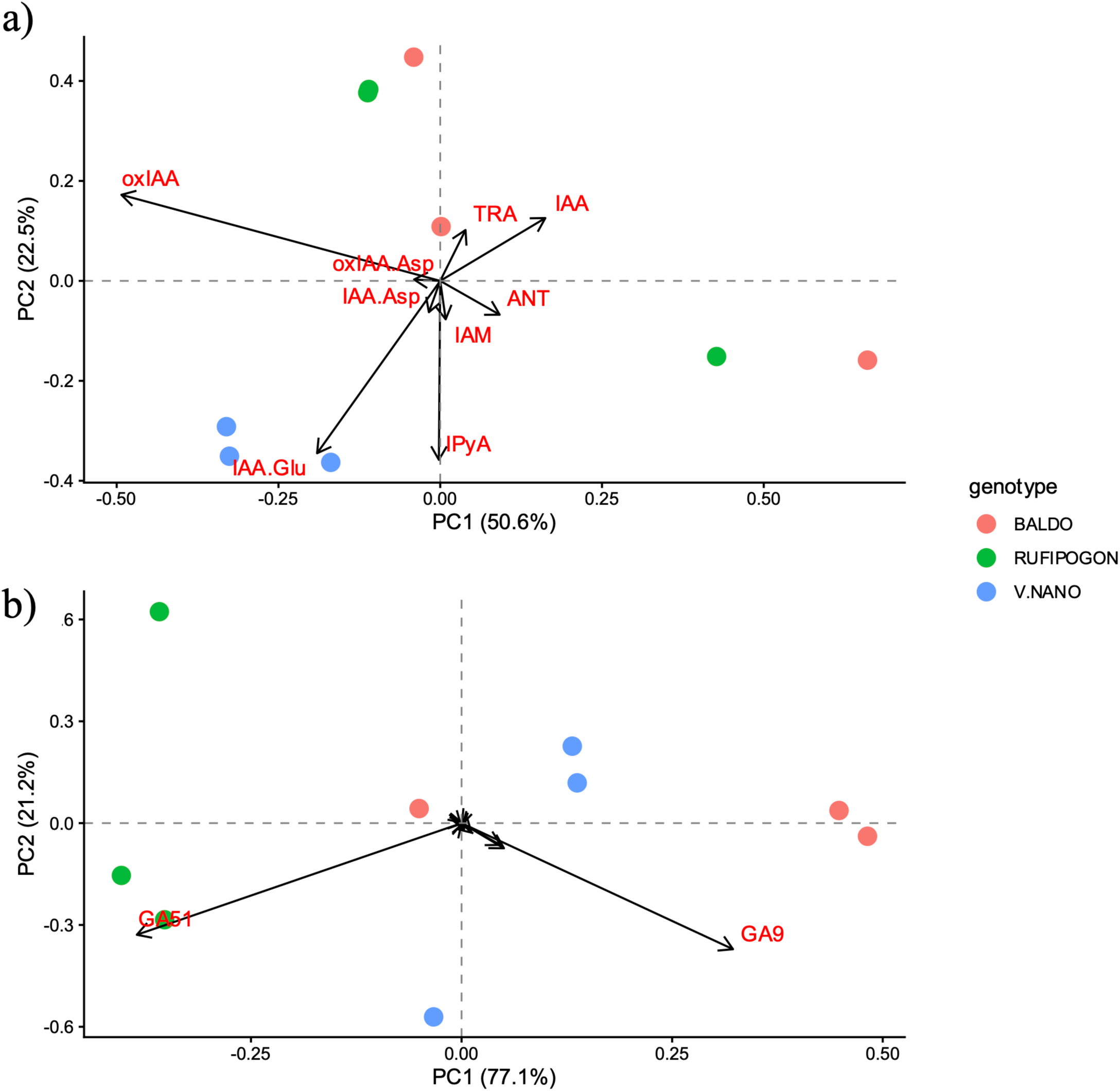
Principal Component Analysis biplots of A) auxins and B) gibberellins profiles in root exudates of the three genotypes. Dots represent biological replicates, while vectors represent metabolite loadings. The percentage of variance explained by PC1 and PC2 is shown on the axes.

### Different host genotypes showed diverse transcriptional responses to the same endophytes

RNA-seq of inoculated rice genotypes (Dataset S2) showed that plants also discriminate between bacterial strains. Gene entries collected from the 54 rice plant samples were filtered to retain genes expressed in at least 3 samples, ensuring consistency across sample replicates. Considering all three plant genotypes (“Overall”), RCA24 affected the expression of more genes than RCA25 (Table 3 and Table S5). When the three plant genotypes were analyzed separately, we observed the fewest number of DEGs for *O. rufipogon* (“All”) vs the control. The stem of the “Baldo” rice showed a distinct response to RCA25 and RCA24, with a much higher number of DEGs (3,813) compared to “Vialone Nano” and *O. rufipogon*. The three plant genotypes differ in their ability to perceive endophytic bacteria colonization: *O. rufipogon* roots showed the greatest transcriptional response to RCA25 inoculation (Table 3 and Table S5), consistent with a stronger activation of this strain by *O. rufipogon* root exudates (Table 2).

**Table 3.** Number of DEGs identified in *O. sativa* and *O. rufipogon*.

|  | Contrast | N. of DEGs | Site |
| --- | --- | --- | --- |
| Overall | RCA24 vs Control | 159 | All |
|  | RCA25 vs Control | 74 | All |
|  | RCA25 vs RCA24 | 93 | All |
| *O. sativa* “Baldo” | RCA25 vs RCA24 | 117 | All |
|  | RCA25 vs RCA24 | 42 | Root |
|  | RCA25 vs RCA24 | 3,813 | Stem |
| *O. sativa* “Vialone Nano” | RCA25 vs RCA24 | 79 | All |
|  | RCA25 vs RCA24 | 102 | Root |
|  | RCA25 vs RCA24 | 77 | Stem |
| *O. rufipogon* | RCA25 vs RCA24 | 39 | All |
|  | RCA25 vs RCA24 | 347 | Root |
|  | RCA25 vs RCA24 | 61 | Stem |

Comparisons with control (blank) and between RCA25 and RCA24 are reported for the whole

The principal component analysis reported in Fig. 5 shows distinct responses for stem and roots and for plant genotypes. There is a strong separation along the first principal component for plant tissues, and along the second principal component for *O. rufipogon* and *O. sativa*. A nested likelihood ratio test (LRT) was also applied to the entire dataset and to roots and shoots separately to analyze tissue-specific effects. Models with increasing complexity were compared to assess the significance of each factor: bacterial strain, plant genotype, and their interaction. For genes whose expression was significantly influenced, plant genotype emerged as the major factor contributing to expression variability, affecting 5.78% of genes in the overall dataset (Table 4). This effect was even more evident in roots (9.80%) and shoots (6.39%). Fewer genes were significantly influenced by the bacterial strains (0.10% overall, 0.05% in roots, and 1.46% in shoots), suggesting a moderate transcriptional impact of strain identity. A small but biologically relevant subset of genes exhibited a significant strain × genotype interaction (0.01% in the full dataset, 0.63% in roots, and 1.55% in shoots), suggesting that specific combinations of rice genotypes and bacterial strains produced distinct transcriptional outcomes. The number of DEGs associated with the interaction was higher when the root (764 DEGs, 0.63% of total number of tested genes) and shoot subcompartments (1,883 DEGs, 1.55%) were considered. The functional enrichment (Fig. 6) of differentially expressed genes (DEGs) whose variance was significantly explained by the interaction term showed that the 10 most enriched GO terms were mainly related to plant defense and stress responses, especially hormone-mediated signaling pathways involved in plant immunity. The regulation of defensive response, the defensive response, and the jasmonic acid-mediated signaling pathway were significantly overrepresented. Focusing on the root transcriptome, the enriched GO terms mainly concerned fundamental cellular processes related to protein synthesis and ribosome function, reflecting increased metabolic and growth-related activities in the root tissue. Differentially expressed shoot-specific genes were mainly enriched in GO terms associated with responses to external stimuli, response to biotic stimuli, response to bacteria, and regulatory processes, indicating that shoots were actively involved in signaling and defense mechanisms against environmental challenges and microbial interactions. When the functions (COG categories) of DEGs for *O. rufipogon* and *O. sativa* were compared with RCA25 and RCA24, distinct patterns were observed (Fig. 7, Fig. 8, Table S1). For ‘Vialone Nano’ and *O. rufipogon*, mainly upregulated genes (RCA25 vs. RCA24) were observed, while for ‘Baldo’, several downregulated genes were also identified. Among the assigned functions, signal transduction mechanisms (COG T) and carbohydrate transport and metabolism (COG G) were the second most common. The largest number of DEGs lacked an assigned COG (COG S) for all three genotypes, suggesting that we are still far from understanding the molecular functions associated with bacterial recognition by plants. Similar findings to those from the COG analysis emerged from the mapping of DEGs to GO categories (Fig. 7, Fig. 8, Table S1), where downregulation of several categories partially related to defense functions was observed, especially in shoot tissue. RCA25 stimulated several GO categories in *O. rufipogon* roots, contrary to the shoot and root of ‘Baldo’ and ‘Vialone Nano’ (Fig. 7, Fig. 8, Table S1).

**Fig. 5.**
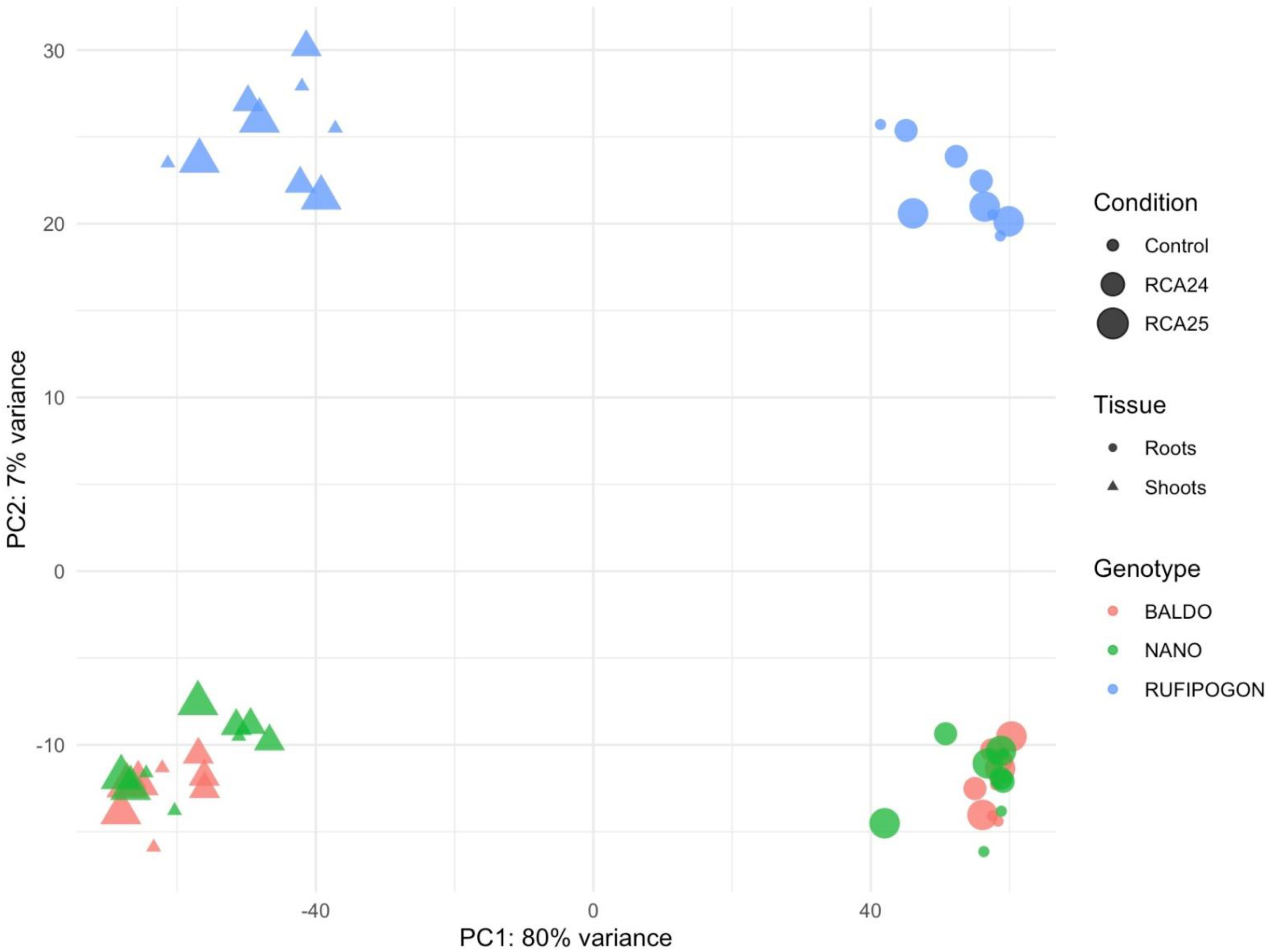
Principal Component Analysis of plant transcriptomes. The percentages of variance for the first and second components are reported on the two axes.

**Fig. 6.**
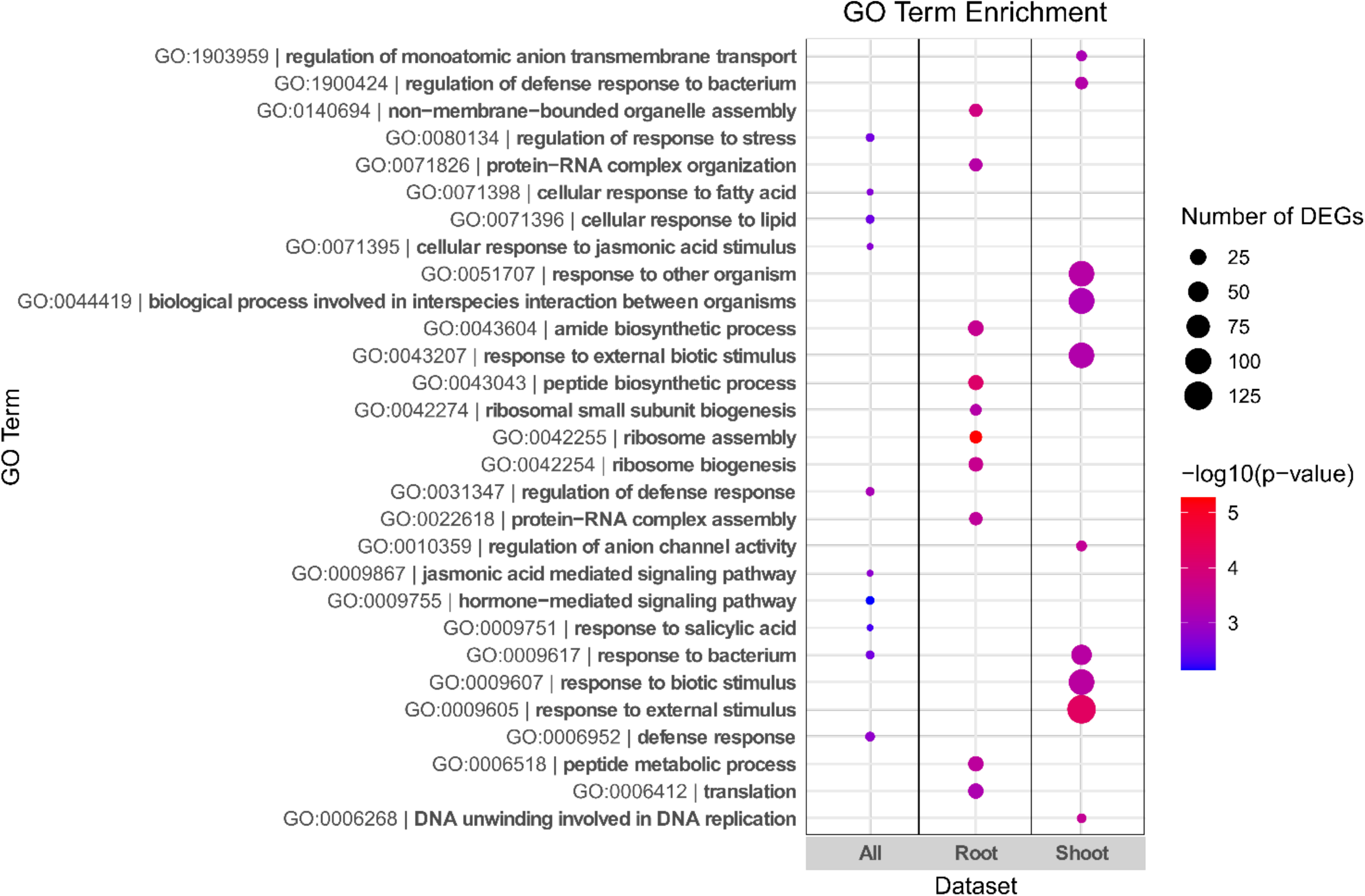
GO enrichment analysis of differentially expressed genes (DEGs) in rice explained by the interaction term in the nested model. The x-axis shows the three datasets considered, while the y-axis displays the enriched biological processes. Bubble size indicates the number of DEGs associated with each GO term for each dataset, and bubble color reflects the statistical significance of enrichment (with darker shades representing lower p-values). For each dataset, the top 10 most significantly enriched GO terms were selected.

**Fig. 7.**
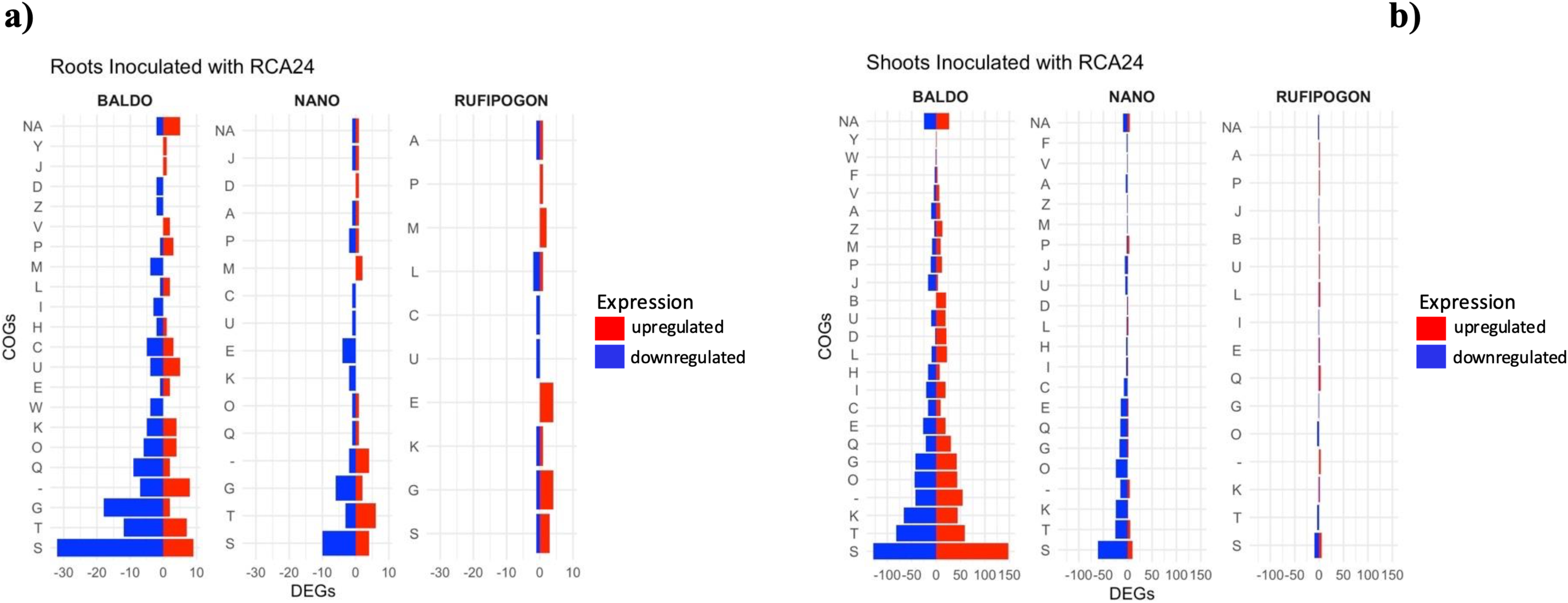
Functional profiling of DEGs obtained by comparing the treatments with RCA24. A) Roots; B) Shoots. Codes indicate: [J] Translation, ribosomal structure and biogenesis; [A] RNA processing and modification; [K] Transcription; [L] Replication, recombination and repair; [B] Chromatin structure and dynamics;[J] Translation, ribosomal structure and biogenesis; [D] Cell cycle control, cell division, chromosome partitioning; [Y] Nuclear structure; [V] Defense mechanisms; [T] Signal transduction mechanisms; [M] Cell wall/membrane/envelope biogenesis; [N] Cell motility; [Z] Cytoskeleton; [W] Extracellular structures; [U] Intracellular trafficking, secretion, and vesicular transport; [O] Posttranslational modification, protein turnover, chaperones; [C] Energy production and conversion; [G] Carbohydrate transport and metabolism; [E] Amino acid transport and metabolism; [F] Nucleotide transport and metabolism; [H] Coenzyme transport and metabolism; [I] Lipid transport and metabolism; [P] Inorganic ion transport and metabolism; [Q] Secondary metabolites biosynthesis, transport and catabolism; [R] General function prediction only; [S] Function unknown.

**Fig. 8.**
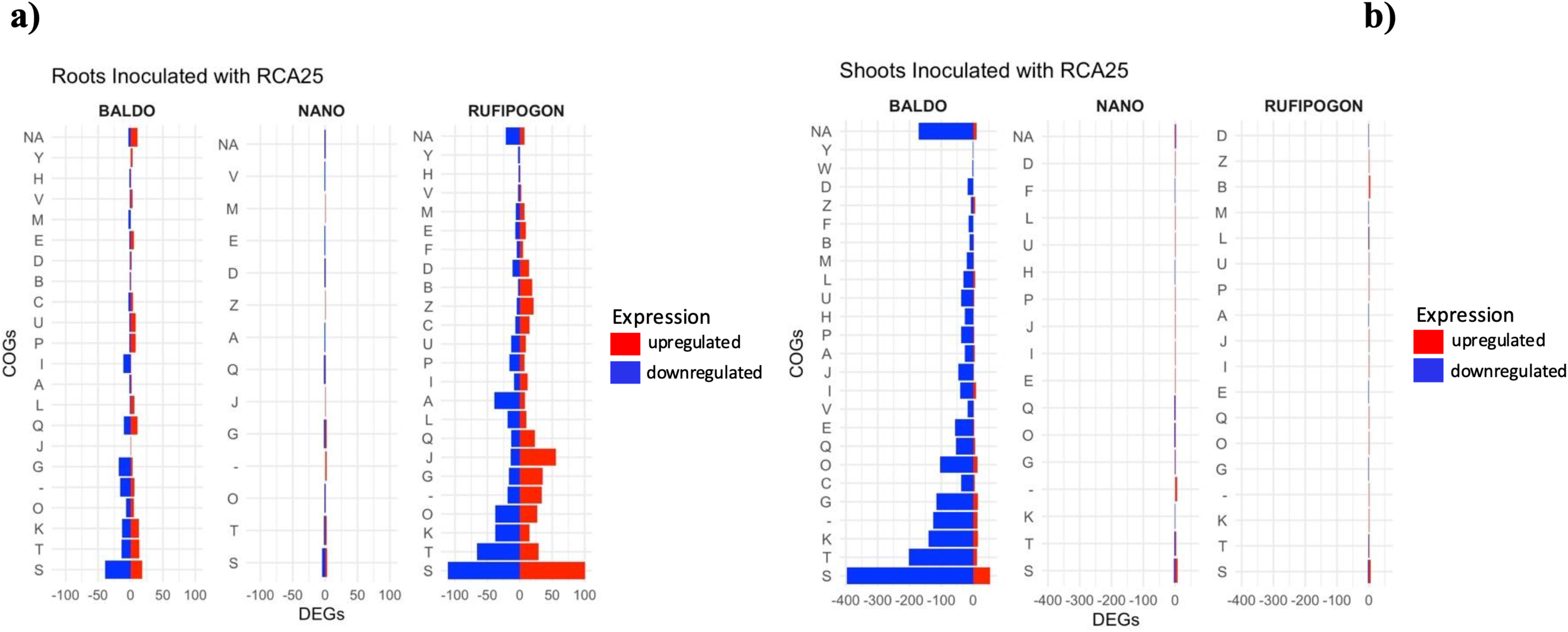
Functional profiling of DEGs obtained by comparing the treatments with RCA25. A) Roots; B) Shoots. Codes indicate: [J] Translation, ribosomal structure and biogenesis; [A] RNA processing and modification; [K] Transcription; [L] Replication, recombination and repair; [B] Chromatin structure and dynamics;[J] Translation, ribosomal structure and biogenesis; [D] Cell cycle control, cell division, chromosome partitioning; [Y] Nuclear structure; [V] Defense mechanisms; [T] Signal transduction mechanisms; [M] Cell wall/membrane/envelope biogenesis; [N] Cell motility; [Z] Cytoskeleton; [W] Extracellular structures; [U] Intracellular trafficking, secretion, and vesicular transport; [O] Posttranslational modification, protein turnover, chaperones; [C] Energy production and conversion; [G] Carbohydrate transport and metabolism; [E] Amino acid transport and metabolism; [F] Nucleotide transport and metabolism; [H] Coenzyme transport and metabolism; [I] Lipid transport and metabolism; [P] Inorganic ion transport and metabolism; [Q] Secondary metabolites biosynthesis, transport and catabolism; [R] General function prediction only; [S] Function unknown.

**Table 4.** Relevance of plant transcriptome in bacterial strain recognition. Summary of the number and percentage of significant gene combinations out of the total 121,464 (40.488 genes) identified through a nested likelihood ratio test (LRT).

| Strain | Number of DEGs<sup>a</sup> |  |  |
| --- | --- | --- | --- |
|  | a) All | b) Root | c) Shoot |
| Rice genotype | 127 | 61 | 1793 |
| Strain and rice | 7003 | 11859 | 7750 |
| Strain x rice genotype | 7130 | 11920 | 9543 |
| None | 12 | 766 | 1865 |
| Total | 84498 | 69135 | 77409 |
| Strain | 105660 (100%) | 105660 (100%) | 105660 (100%) |

<sup>a</sup>Significance was determined using an FDR-adjusted p-value threshold of 0.05. Models were applied to the a) full transcriptome dataset as well as to b) root and c) shoot subsets. Genes were categorized based on the microbial strain (i.e., differentially expressed only between strains), by the rice genotype (i.e., differentially expressed only between rice genotypes), with both factors independently (i.e., differentially expressed in relation to both rice genotype and microbial strain, but not explained by their interaction), or with the interaction between rice genotype and microbial strain (i.e., expression significantly influenced by the specific combination of genotype and strain).

## Discussion

Understanding the molecular mechanisms underlying the interaction between plants and their associated microbial communities is crucial for developing sustainable and efficient agricultural practices (Ke *et al*., 2021; Busby *et al*., 2017).

Here, we have provided molecular data supporting species-specific colonization of rice plants by distinct bacterial endophytes. The observation that distinct rice genotypes produce different phytohormones cocktails in their root exudates and, consequently, elicit variable transcriptional responses in bacterial strains highlights the importance of genotype-by-genotype interactions in shaping plant-microbe associations and possible phytohormone-mediated communication between plants and bacteria. Our results are consistent with previous findings in rhizobia-legume systems, where plant genotype-driven differences in overall root exudate metabolites have been shown to influence the expression of microbial genes crucial for symbiosis formation (Fagorzi *et al*., 2021; Kiers *et al*., 2007). When the bacterial transcriptomes were analyzed in the presence of plant root exudates, the differential transcriptional responses in RCA24 and RCA25 suggested that bacterial genotype is a key determinant of the interaction and partially reflects colonization titers. For instance, RCA25 exhibited a stronger response to root exudates of *O. rufipogon* than those of *O. sativa* accessions, suggesting that RCA25 may be better adapted to this ancestral rice genotype. In contrast, RCA24 showed a marked ability to distinguish between the *O. sativa* varieties, *O. rufipogon* and “Vialone Nano”, with the latter eliciting the highest percentage of unique DEGs, thus indicating its ability to adapt to different host environments. Future metabolomic analyses and fractionations of root exudate metabolomes from different rice varieties could help identify potential metabolites responsible for the marked differences in RCA24 transcriptomic response across rice varieties. It could be hypothesized that the differential transcriptional responses observed in the early stages of plant-microbe interactions contribute to subsequent colonization patterns and plant phenotypes. However, we must consider that we are observing a transcriptional response to root exudates under gnotobiotic conditions *in vitro*, which may not reflect the conditions experiences by these strains when they approach roots and plants in natural soil. Our findings highlighted that *O. rufipogon* root exudates possess distinct chemical signatures that strongly modulate bacterial behavior, suggesting that the domestication process may have inadvertently altered the chemical composition of root exudates, thereby influencing the recruitment and activity of beneficial microbes. The analysis of phytohormone composition in the root exudates of *O. rufipogon* and *O. sativa* varieties revealed marked differences in the gibberellin GA_9_ and GA_51_. The biological significance of these differences in phytohormone composition with respect to transcriptomic response of RCA25 (which discriminated against *O. rufipogon*) and RCA24 (which discriminated against *O. sativa* varieties) is beyond the aim of the present work. However, it is worth noting that bacteria possess a cytochrome P450-based pathway for gibberellin biosynthesis (Salazar-Cerezo *et al*., 2018) and that gibberellin biosynthesis has been reported in the rice leaf streak pathogen *Xanthomonas oryzae* pv. *oryzicola* (Nagel *et al*., 2017). Additionally, IAA was present at concentrations higher in *O. sativa* varieties than in *O. rufipogon*. IAA was produced at fairly high levels only by RCA24 (Andreozzi *et al*., 2019). It is known that bacteria can also sense IAA, which regulates biofilm formation, metabolism, auxin catabolism, virulence, chemotactic responses, and host plant colonization (Etesami and Glick, 2024; Rico-Jiménez *et al*., 2024). Indeed, it has been demonstrated that IAA produced by a variety of endophytic bacteria plays a key role in establishing beneficial relationships between plants and microorganisms, as it helps them manipulate plant physiology to their advantage despite IAA is use also by pathogenic bacteria to overcome plant defense systems (Arora *et al*., 2024; Feng *et al*., 2024). In *Serratia plymuthica* A153, as observed in this study for RCA25, IAA induces a strong up-regulation of genes assigned to cell wall/membrane/envelope biogenesis (COG M), as well as amino acid transport and metabolism (COG E) (Rico-Jiménez *et al*., 2024). Therefore, we cannot *a priori* rule out that *K. sacchari* RCA25 and *E. asburiae* RCA24 transcriptomes may be influenced by the different phytohormone composition of root exudates. These findings are consistent with previous reports on other grass species (e.g., barley), for which domestication has been shown to alter the composition and complexity of root exudates, thereby reducing interactions with beneficial microbes (Escudero-Martinez and Bulgarelli, 2019). Considering the specific functional features of DEGs, we also observed that perception of plant root exudates induced a general remodeling of energy fluxes inside the cell. A general remodeling of such central metabolic pathways was also reported for the legume symbiont *Sinorhizobium meliloti* (DiCenzo *et al*., 2016; Fagorzi *et al*., 2021). Plant genes shaping the microbiota composition have been recently identified in barley, tomato, and sorghum (Wu *et al*., 2012; Deng *et al*., 2021; Escudero-Martinez *et al*., 2022; Oyserman *et al*., 2022). In rice, it was observed that soil microbiome triggers the downregulation of genes coding for proteins with an NB-ARC domain (leucine-rich repeat receptor-like kinases and nucleotide-binding leucine-rich repeat receptors) in roots (Santos-Medellin *et al*., 2022). We hypothesize that genes involved in defense-related pathways may contribute to the transcriptomic differences observed between *O. rufipogon* and *O. sativa*. Therefore, we performed a transcriptomic experiment with rice plants of *O. rufipogon, O. sativa* “Baldo”, and “Vialone Nano” subjected to colonization by RCA25 or RCA24. Differences in the number of DEGs observed in this analysis highlighted that RCA24 preferentially modulated gene expression in ‘Vialone Nano’, while RCA25 preferentially modulated gene expression in *O. rufipogon* and ‘Baldo’. This difference reflected changes in the expression of genes actively involved in endophytic colonization.

Rice transcriptome results revealed that while plant genotype was the dominant factor in shaping transcriptional responses to bacterial colonization, strain × genotype interactions affected a biologically significant subset of genes. The hypothesis of plant transcriptome signatures of bacterial host preference was supported by the highest number of plant’s DEGs observed in *O. rufipogon* treated with RCA25 and Baldo treated with RCA24. The nested-LRT model showed that the shoot transcriptome displayed stronger interaction effects than the root transcriptome, suggesting that systemic signaling and aboveground perception of bacterial presence may play a key role in determining colonization outcomes. We then speculated that rice’s response to bacterial colonization occurred primarily when the bacteria reached the stem tissue, where differences in colonization titers were indeed observed. The genes for which the transcriptome of RCA24 and RCA25 differed most were involved in plant defense (e.g., cystatin genes) and hormone-mediated signaling, supporting the idea that plants actively discriminate between endophytic strains, likely balancing growth promotion and immune surveillance. These findings confirmed those obtained in barley and tomato (Escudero-Martinez *et al*., 2022; Oyserman *et al*., 2022), where plant defence regulators such as NPR1 have been shown to underlie microbiota assembly, indicating that conserved defence-related genetic modules may influence microbial recruitment across species. Interestingly, the magnitude and specificity of the plant response differed among rice genotypes. The cultivated variety Baldo displayed a particularly high number of DEGs in stems when compared with RCA24 and RCA25 colonization, whereas the wild ancestor *O. rufipogon* exhibited fewer transcriptional changes in this tissue. This result suggests that domestication may have altered rice’s transcriptional plasticity towards endophytes, potentially making the two modern cultivars analyzed more responsive, resulting in less effective colonization. Conversely, a significant number of DEGs were observed in roots colonized by RCA25, suggesting that elective recognition processes are involved in this interaction. However, it should be noted that a high fraction of DEGs lacked functional annotation, indicating that several factors in both plant and bacterial genomes remain to be elucidated to understand microbe-plant interaction fully.

Our data were obtained with a simplified system (three rice varieties and two bacteria isolated from the same cultivated rice field), compared with natural plant-microbiota interactions, which involve a single plant interacting with hundreds of different microorganisms and with the root exudates of surrounding plants. Nonetheless, they revealed considerable variability in the response of both the plants and their different compartments, as well as the bacteria.

The results presented in this work highlight the immense complexity of interactions between individual plants and their microbiota, a complexity that has so far been largely underestimated. Further studies integrating metabolomics with plant and bacterial transcriptomes could clarify whether hormone composition directly triggers transcriptional changes in plants or whether these changes are indirectly influenced by bacterial metabolism.

Given that crop plants have adapted to new growing conditions that preclude some of the capabilities conserved by more resistant or ancestral plants, our findings pave the way for a recovery of both microbial and plant biodiversity, promising significant advances in knowledge and potential applications to support plant development under changing climate conditions, potentially even restoring traits lost during domestication (Raaijmakers & Kiers, 2022).

## Conclusions

Overall, these data highlight that rice plants not only release genotype-specific molecular signals but also develop distinct transcriptional programs in response to endophytic colonization. The bacterial transcriptional response to ancestral rice root exudates, such as *O. rufipogon*, reported in this study highlight that bacteria communicate *via* exudates with the host plants by activating specific gene pathways in their roots. Future studies integrating metabolomics with plant and bacterial transcriptomes could clarify whether hormone composition directly triggers transcriptional changes in plants or whether these are indirectly shaped by bacterial metabolism. A deeper understanding of the genetic basis of these changes could inform breeding programs to develop rice cultivars with enhanced microbial associations and improved agronomic traits.

## Supplementary Information

The following supplementary materials are available at PCR online.

**Table S1.** Description of single-letter COG codes.

**Table S2.** Mean concentrations (pmol/l ± SD, n = 3) of auxin in root exudates and extracts of *O. sativa* cv. Baldo, *O. rufipogon*, and *O. sativa* cv. Vialone Nano. Values below the limit of detection are reported as the LOD threshold and highlighted in bold.

**Table S3.** Mean concentrations (pmol/l ± SD, n = 3) of gibberellins in root exudates and extracts of *O. sativa* cv. Baldo, *O. rufipogon*, and *O. sativa* cv. Vialone Nano. Values below the limit of detection are reported as the LOD threshold and highlighted in bold.

**Table S4.** Description of auxins codes.

**Table S4.** Number of DEGs identified in *O. sativa* and *O. rufipogon* for all analyzed contrasts.

**Fig. S1.** Principal Component Analysis (PCA) biplots of A) auxins and B) gibberellins profiles in root extracts of the three genotypes. Dots represent biological replicates, while vectors represent metabolite loadings. The percentage of variance explained by PC1 and PC2 is shown on the axes.

**Dataset S1.** Details of metabolic pathways (KEGG mapped) of differentially expressed genes identified for *E. asburiae* RCA24 and *K. sacchari* RCA25 treated with rice root exudates.

**Dataset S2.** List of differentially expressed genes identified by comparing transcription profiles of plants colonized with *E. asburiae* RCA24 and *K. sacchari* RCA25 and uninoculated control ones.

## Acknowledgments

The authors gratefully acknowledge Marco Petruzziello, Francesca Segreti, and Antonio Suppa for their technical support. We also thank Magdalena Vlčková and Andrea Novotná for technical assistance.

## Author contributions

Conceptualization: CB and AM; Methodology: FV, IP, CF, MLA, AR, SV, DT, AP, IV, FB; Data Analysis: FV, IP, CF, and MLA; Writing (review & editing): FV, CF, VB, AR, GV, EM, RD, AM, CB; Supervision: CB and AM; Funding: CB, VB and AM

## Data availability

Sequence reads of RNA-seq have been deposited and are available as Projects PRJEB82954 (ENA database, bacteria RNA-Seq data) and PRJNA1309287 (NCBI database, plant RNA-Seq data).

All the custom code used for pre-processing and data analysis is available at the following GitHub link: https://github.com/IacopoPasseri/RIS8-Ricebiota.

## Declarations

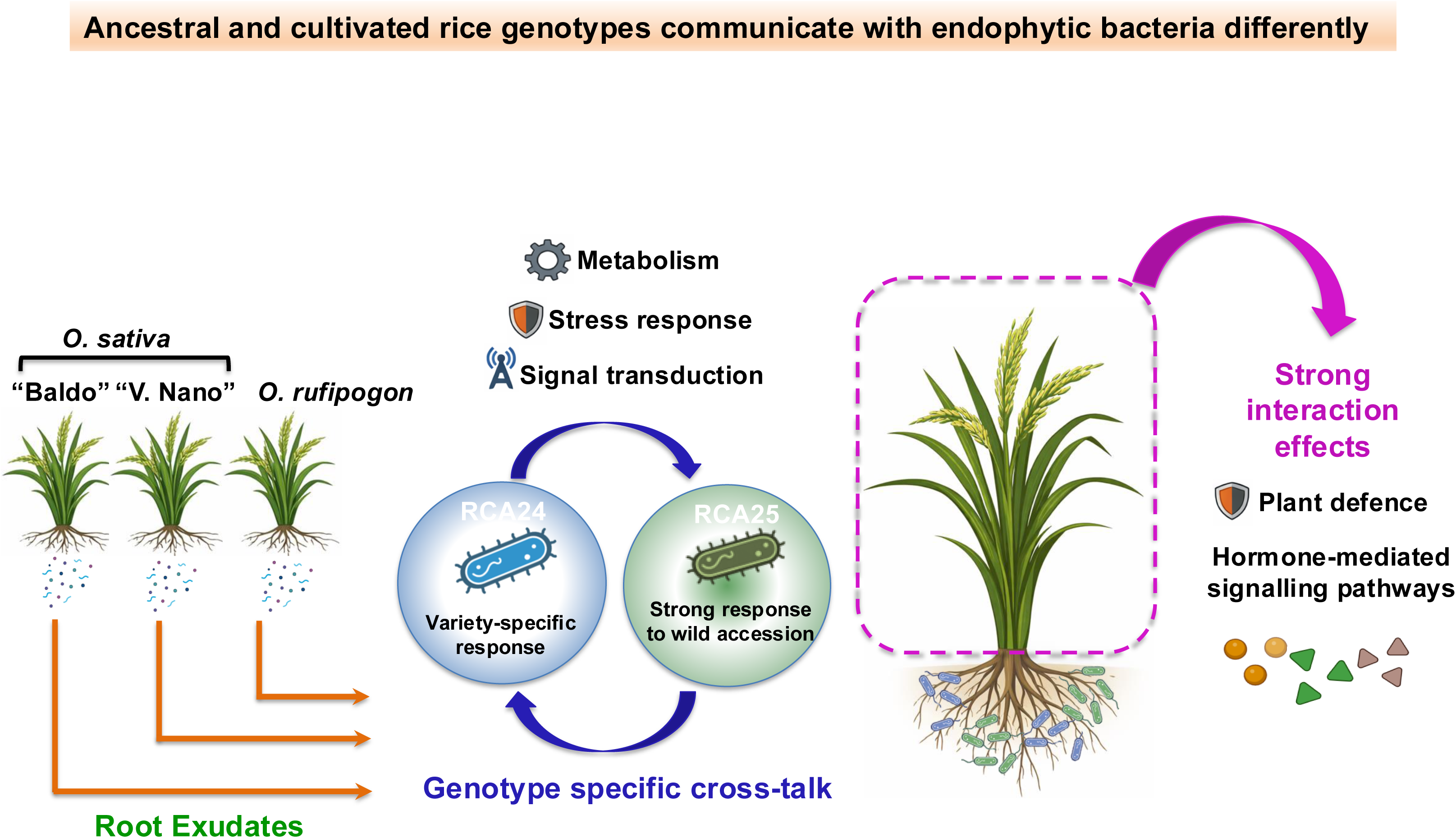

