## Supplementary material for "Wild rice *Oryza rufipogon* outperforms cultivated rice in stimulating beneficial bacterial endophytes": Supplental Tables and Figures

**Supplementary Tables and Figures**

**Table S1**. Description of single-letter COG codes.

| |
|---|
| INFORMATION STORAGE AND PROCESSING |
| [J] Translation, ribosomal structure and biogenesis<br>[A] RNA processing and modification<br>[K] Transcription<br>[L] Replication, recombination and repair<br>[B] Chromatin structure and dynamics |
| CELLULAR PROCESSES AND SIGNALING |
| [D] Cell cycle control, cell division, chromosome partitioning<br>[Y] Nuclear structure<br>[V] Defense mechanisms<br>[T] Signal transduction mechanisms<br>[M] Cell wall/membrane/envelope biogenesis<br>[N] Cell motility<br>[Z] Cytoskeleton<br>[W] Extracellular structures<br>[U] Intracellular trafficking, secretion, and vesicular transport<br>[O] Posttranslational modification, protein turnover, chaperones |
| METABOLISM |
| [C] Energy production and conversion<br>[G] Carbohydrate transport and metabolism<br>[E] Amino acid transport and metabolism<br>[F] Nucleotide transport and metabolism<br>[H] Coenzyme transport and metabolism<br>[I] Lipid transport and metabolism<br>[P] Inorganic ion transport and metabolism<br>[Q] Secondary metabolites biosynthesis, transport and catabolism |
| POORLY CHARACTERIZED |
| [R] General function prediction only<br>[S] Function unknown |

**Table S2**. Mean concentrations (pmol/l ± SD, n = 3) of *auxins* in root exudates of *O. sativa* cv. Baldo, *O. rufipogon*, and *O. sativa* cv. Vialone Nano. Values below the limit of detection are reported as the LOD threshold and highlighted in bold.

| Compound | | Baldo | Rufipogon | V. Nano |
|---|---|---|---|---|
| ANT | exudate | 45.4333 ± 49.7968 | 16.3667 ± 8.3339 | 26.0000 ± 13.6963 |
| | root extract | 495.9333 ± 91.1114 | 875.5333 ± 334.3919 | 2243.0000 ± 1245.7953 |
| IAA | exudate | 33.2000 ± 30.6886 | 20.2667 ± 7.4849 | 10.7000 ± 4.6893 |
| | root extract | 1177.8667 ± 73.8419 | 411.0333 ± 76.4690 | 4840.9333 ± 3103.4373 |
| IAA-Asp | exudate | 0.4767 ± 0.4210 | 0.3202 ± 0.2770 | 0.8367 ± 0.3609 |
| | root extract | 678.6233 ± 98.9018 | 125.0133 ± 124.1091 | 1506.3567 ± 639.0733 |
| IAA-Glu | exudate | **0.0050 ± 0.0000** | **0.0050 ± 0.0000** | 4.0000 ± 0.2646 |
| | root extract | 425.6333 ± 88.1104 | 35.2000 ± 10.0583 | 842.3333 ± 67.7521 |
| IAA-glc | exudate | **0.0050 ± 0.0000** | **0.0050 ± 0.0000** | **0.0050 ± 0.0000** |
| | root extract | **0.0050 ± 0.0000** | 24.3667 ± 7.9103 | 15.3683 ± 14.6576 |
| IAM | exudate | 0.6767 ± 0.6199 | 0.2633 ± 0.1686 | 0.8800 ± 0.4838 |
| | root extract | 26.0700 ± 6.5608 | 15.6800 ± 3.0086 | 28.6333 ± 5.9057 |
| IAN | exudate | **0.0100 ± 0.0000** | **0.0100 ± 0.0000** | **0.0100 ± 0.0000** |
| | root extract | **0.0100 ± 0.0000** | **0.0100 ± 0.0000** | **0.0100 ± 0.0000** |
| IPyA | exudate | 179.1667 ± 89.8255 | 158.2000 ± 103.3561 | 392.0000 ± 154.3210 |
| | root extract | 30795.0000 ± 1480.8507 | 13627.9000 ± 5074.4723 | 49191.8000 ± 21702.7721 |
| TRA | exudate | 2.3233 ± 2.1495 | 0.8200 ± 0.0265 | 0.8333 ± 0.8702 |
| | root extract | 381.3667 ± 124.1259 | 298.4367 ± 74.9507 | 425.3967 ± 174.6320 |
| TRP | exudate | **0.0100 ± 0.0000** | **0.0100 ± 0.0000** | **0.0100 ± 0.0000** |
| | root extract | 1176752.2667 ± 175454.2987 | 1193205.7000 ± 202279.8735 | 1656015.3333 ± 575500.8945 |
| oxIAA | exudate | 24.4000 ± 23.7956 | 41.8000 ± 35.0801 | 61.4333 ± 18.0059 |
| | root extract | 206.5667 ± 72.0723 | 72.7333 ± 16.9223 | 651.0333 ± 123.2742 |
| oxIAA-Asp | exudate | 0.1137 ± 0.1470 | 0.5567 ± 0.4535 | 0.4633 ± 0.2650 |
| | root extract | 88.4500 ± 8.4143 | 79.4967 ± 62.8399 | 376.6667 ± 288.4398 |
| oxIAA-Glu | exudate | **0.0050 ± 0.0000** | **0.0050 ± 0.0000** | **0.0050 ± 0.0000** |
| | root extract | **0.0005 ± 0.0000** | **0.0005 ± 0.0000** | 13.0333 ± 5.8535 |
| oxIAA-glc | exudate | **0.0050 ± 0.0000** | **0.0050 ± 0.0000** | **0.0050 ± 0.0000** |
| | root extract | 83.3000 ± 15.2148 | 27.0000 ± 11.9528 | 83.2000 ± 13.0817 |

**Table S3**. Mean concentrations (pmol/l ± SD, n = 3) of *gibberellins* in root exudates of *O. sativa* cv. Baldo, *O. rufipogon*, and *O. sativa* cv. Vialone Nano. Values below the limit of detection are reported as the LOD threshold and highlighted in bold.

| Compound | | Baldo | Rufipogon | V. Nano |
|---|---|---|---|---|
| GA1 | exudate | 0.4167 ± 0.0404 | 0.4233 ± 0.0321 | 0.8233 ± 0.2511 |
| | root extract | 10.3767 ± 4.0501 | 4.1567 ± 0.2301 | 4.8467 ± 0.9136 |
| GA15 | exudate | 0.0043 ± 0.0000 | 0.0043 ± 0.0000 | 0.0043 ± 0.0000 |
| | root extract | **0.0043 ± 0.0000** | **0.0043 ± 0.0000** | **0.0043 ± 0.0000** |
| GA19 | exudate | 0.0773 ± 0.0170 | 0.0863 ± 0.0342 | 0.1140 ± 0.0250 |
| | root extract | 1.0660 ± 0.3461 | 0.6597 ± 0.1007 | 0.8350 ± 0.2114 |
| GA20 | exudate | 3.7533 ± 0.8580 | 1.5433 ± 1.0782 | 3.0567 ± 0.1779 |
| | root extract | 1.2900 ± 0.1345 | 1.5833 ± 0.4744 | 1.2067 ± 0.3252 |
| GA24 | exudate | **0.0095 ± 0.0000** | **0.0095 ± 0.0000** | **0.0095 ± 0.0000** |
| | root extract | **0.0095 ± 0.0000** | **0.0095 ± 0.0000** | **0.0095 ± 0.0000** |
| GA29 | exudate | 0.1133 ± 0.0493 | 0.0800 ± 0.0361 | 0.1867 ± 0.0451 |
| | root extract | 1.1733 ± 0.2350 | 0.7700 ± 0.3041 | 0.4633 ± 0.2616 |
| GA3 | exudate | 1.7167 ± 0.0289 | 1.6500 ± 0.0600 | 1.6600 ± 0.0300 |
| | root extract | 7.1533 ± 0.2403 | 6.7567 ± 0.2237 | 7.4000 ± 0.0265 |
| GA34 | exudate | 0.0077 ± 0.0021 | 0.0100 ± 0.0053 | 0.0433 ± 0.0058 |
| | root extract | **0.0058 ± 0.0000** | **0.0058 ± 0.0000** | 0.0913 ± 0.0696 |
| GA4 | exudate | 1.1367 ± 0.5262 | 0.3100 ± 0.1179 | 2.8167 ± 1.1951 |
| | root extract | 3.2300 ± 2.6601 | 1.2567 ± 0.2201 | 6.3700 ± 6.5314 |
| GA44 | exudate | 0.1833 ± 0.1474 | 0.0186 ± 0.0187 | 0.0057 ± 0.0000 |
| | root extract | 1.5800 ± 0.8643 | 1.3467 ± 0.8658 | 0.7352 ± 0.6450 |
| GA51 | exudate | 11.2367 ± 19.4538 | 577.4333 ± 459.0523 | 74.8333 ± 120.7850 |
| | root extract | **0.0051 ± 0.0000** | **0.0051 ± 0.0000** | 23.3000 ± 36.2352 |
| GA53 | exudate | 83.2000 ± 10.6691 | 66.9000 ± 10.8917 | 84.2667 ± 31.0536 |
| | root extract | 95.0000 ± 11.5295 | 95.7333 ± 8.9668 | 122.4000 ± 67.6610 |
| GA6 | exudate | 0.0087 ± 0.0006 | 0.0090 ± 0.0010 | 0.0083 ± 0.0021 |
| | root extract | 0.0213 ± 0.0023 | 0.0230 ± 0.0130 | 0.0171 ± 0.0089 |
| GA7 | exudate | 0.0433 ± 0.0058 | 0.0300 ± 0.0100 | 0.0367 ± 0.0058 |
| | root extract | 0.3500 ± 0.0458 | 0.3700 ± 0.0436 | 0.2800 ± 0.0436 |
| GA8 | exudate | 0.1767 ± 0.0635 | 0.1333 ± 0.0551 | 0.2400 ± 0.0624 |
| | root extract | 0.7300 ± 0.1400 | 0.8300 ± 0.0964 | 0.7133 ± 0.1415 |
| GA9 | exudate | 956.4333 ± 856.1411 | 13.2333 ± 14.1797 | 219.2000 ± 225.8027 |
| | root extract | 2526.5333 ± 2777.7548 | 687.8667 ± 962.6672 | 22947.6667 ± 36851.4079 |

**Table S4**. Description of auxins codes.

| Compound name | Abbreviation |
|---|---|
| Anthranilic acid | ANT |
| Tryptophan | TRP |
| Tryptamine | TRA |
| Indole-3-acetamid | IAM |
| Indole-3-acetonitrile | IAN |
| Indole-3-pyruvic acid | IPyA |
| Indole-3-acetic acid | IAA |
| IAA-aspartate | IAA-Asp |
| IAA-glutamate | IAA-Glu |
| 2-oxoindole-3-acetic acid | oxIAA |
| oxIAA-aspartate | oxIAA-Asp |
| oxIAA-glutamate | oxIAA-Glu |
| IAA-glucose | IAA-glc |
| oxIAA-glucose | oxIAA-glc |

**Table S5**. Number of DEGs identified in *O. sativa* and *O. rufipogon* for all analyzed contrasts.

| | **Contrast** | **n. of DEGs** | **Site** |
|---|---|---|---|
| Overall | RCA24 vs Control | 159 | All |
| | RCA25 vs Control | 74 | All |
| | RCA25 vs RCA24 | 93 | All |
| *O. sativa* "Baldo" | RCA24 vs Control | 165 | All |
| | RCA25 vs Control | 172 | All |
| | RCA25 vs RCA24 | 117 | All |
| | RCA24 vs Control | 168 | Root |
| | RCA25 vs Control | 274 | Root |
| | RCA25 vs RCA24 | 42 | Root |
| | RCA24 vs Control | 1202 | Stem |
| | RCA25 vs Control | 1870 | Stem |
| | RCA25 vs RCA24 | 3,813 | Stem |
| *O. sativa* "Vialone Nano" | RCA24 vs Control | 92 | All |
| | RCA25 vs Control | 16 | All |
| | RCA25 vs RCA24 | 79 | All |
| | RCA24 vs Control | 51 | Root |
| | RCA25 vs Control | 31 | Root |
| | RCA25 vs RCA24 | 102 | Root |
| | RCA24 vs Control | 274 | Stem |
| | RCA25 vs Control | 45 | Stem |
| | RCA25 vs RCA24 | 77 | Stem |
| *O. rufipogon* | RCA24 vs Control | 35 | All |
| | RCA25 vs Control | 38 | All |
| | RCA25 vs RCA24 | 39 | All |
| | RCA24 vs Control | 24 | Root |
| | RCA25 vs Control | 903 | Root |
| | RCA25 vs RCA24 | 347 | Root |
| | RCA24 vs Control | 46 | Stem |
| | RCA25 vs Control | 33 | Stem |
| | RCA25 vs RCA24 | 61 | Stem |

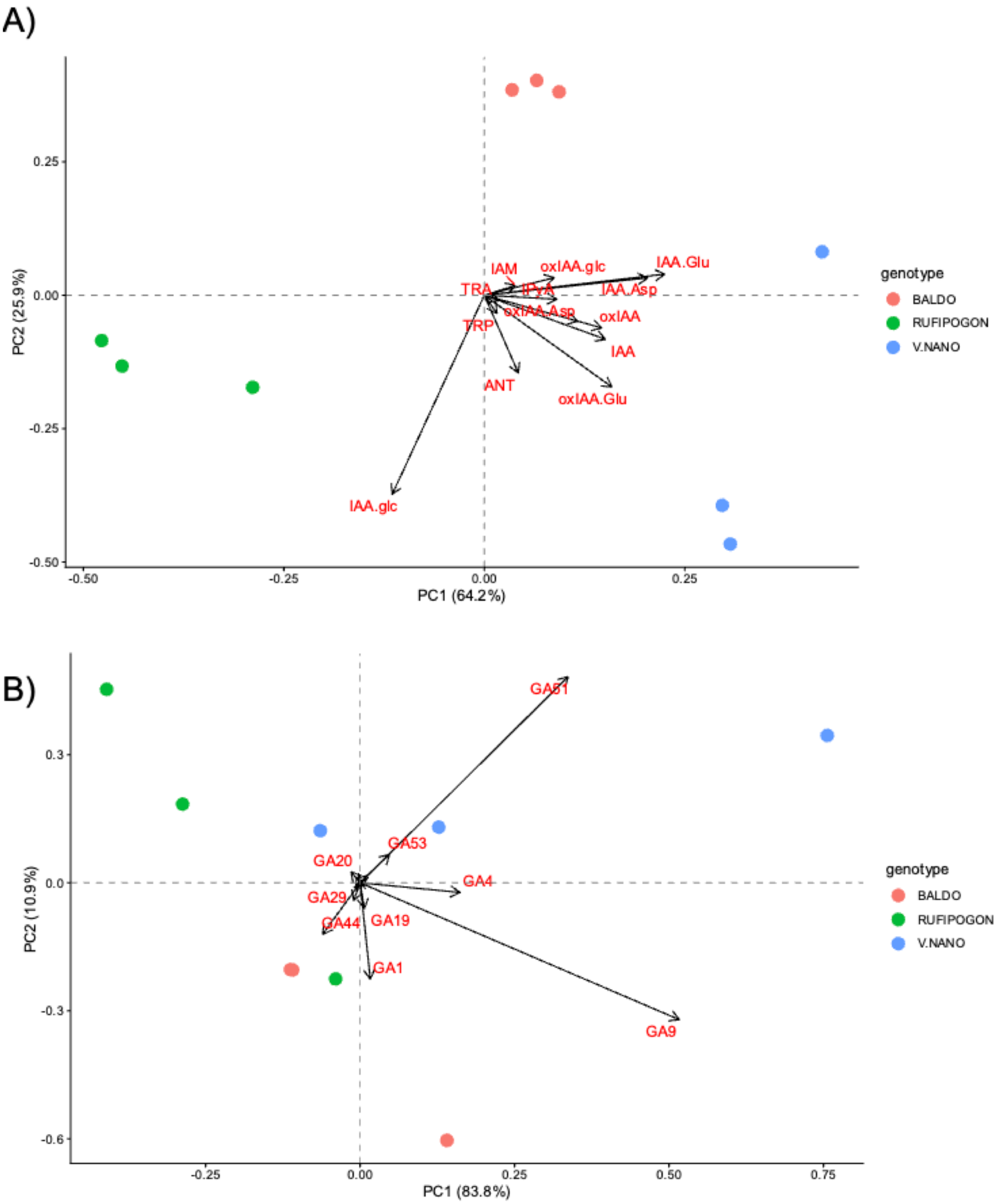


**Fig. S1.** Principal Component Analysis (PCA) biplots of A) auxins and B) gibberellins profiles in root extracts of the three genotypes. Dots represent biological replicates, while vectors represent metabolite loadings. The percentage of variance explained by PC1 and PC2 is shown on the axes.
